# HLAResolver: A statistical framework for resolving HLA ambiguity in T-cell response data

**DOI:** 10.64898/2026.10.29.762841

**Authors:** Hussein Al-Asadi, Matthew T. Noakes, Mark Klinge, Bryan Howie, Harlan Robins

**Affiliations:** Adaptive Biotechnologies, Seattle, Washington, USA

## Abstract

Large-scale models of T-cell recognition require training examples that specify the complete T-cell receptor (TCR)–peptide –human leukocyte antigen (HLA) complex. Multiplexed Identification of T-cell Receptor Antigen Specificity (MIRA) is a high-throughput assay that identifies TCRs associated with responses to specific peptides, but does not determine which HLA presents each peptide. Because a donor can carry up to six distinct class I HLA-A, HLA-B, and HLA-C alleles, each response may have several candidate presenting HLAs.

To narrow this set of candidate presenting HLAs, we developed HLAResolver, a statistical framework that combines MIRA response patterns across HLA-typed donors with NetMHCpan predictions of peptide presentation. For each peptide, HLAResolver uses negative-binomial regression to jointly estimate associations between HLA alleles and TCR response counts, accounting for differences in assay yield. NetMHCpan eluted-ligand predictions guide sparse regularization, favoring alleles with stronger predicted presentation while allowing strong MIRA response evidence to support alternatives.

To assess whether this integration provides an advantage over either approach alone, we compared HLAResolver with NetMHCpan, which predicts peptide–HLA presentation, and the Restrictor Analysis Tool for Epitopes (RATE), which infers HLA associations from T-cell response patterns. Using held-out MIRA experiments, we assessed each method’s ability to predict observed responses and the number of candidate HLAs it assigned. At matched specificity for predicting MIRA responses, HLAResolver achieved higher sensitivity while selecting fewer candidate HLAs per peptide than either comparator.

## Introduction

MIRA identifies peptide-responsive TCRs at high throughput by testing peptide antigens in the context of a donor’s antigen-presenting cells (Klinger et al., 2015). The donor’s HLA genotype therefore defines the possible presenting molecules, but MIRA does not identify the restricting HLA for an individual TCR-peptide response. A donor can carry up to six distinct class I HLA-A, HLA-B, and HLA-C alleles, and multiple alleles may plausibly present the same peptide.

Each MIRA TCR-peptide association therefore corresponds to a set of candidate TCR-peptide-HLA complexes rather than a single complex.

Resolving this uncertainty would make MIRA data more useful for TCR-pMHC modeling. Two existing methods provide different types of evidence. NetMHCpan predicts whether a peptide is likely to be presented by an HLA allele (Jurtz et al., 2017; Reynisson et al., 2020), but permissive thresholds can identify many possible alleles. The Restrictor Analysis Tool for Epitopes (RATE) tests whether carrying an HLA allele is statistically associated with response to a peptide (Paul et al., 2015, 2017). Because RATE tests binary responses one allele at a time, it can be difficult to separate the effects of HLA alleles in linkage disequilibrium (LD).

HLAResolver combines these approaches in a negative-binomial regression model of peptide-specific MIRA TCR counts, adjusting for differences in assay yield across experiments. For each peptide, the model estimates the effects of candidate HLA alleles jointly, helping distinguish associations among alleles in LD. NetMHCpan eluted-ligand percentile ranks (%Rank_EL) guide allele-specific L1 regularization, favoring alleles with stronger predicted presentation while allowing MIRA response evidence to support alternatives. The resulting sparse set of candidate HLAs provides annotations for TCR –peptide associations for downstream modeling and experimental follow-up.

Immunopeptidomics presents a similar problem. Mass spectrometry can identify peptides presented by an HLA-typed sample without showing which of its alleles presents each peptide. Methods such as GibbsCluster and MixMHCp infer allele-associated binding motifs from these multi-allelic peptide data (Bassani-Sternberg and Gfeller, 2016; Andreatta et al., 2017). HLAResolver also addresses uncertain HLA assignments, but uses TCR response patterns across HLA-typed MIRA experiments rather than peptide-binding motifs.

Evaluating HLAResolver is challenging because experimentally confirmed HLA restrictions are not available at the scale of the MIRA dataset. We therefore asked whether HLA assignments learned from training experiments could predict peptide responses in held-out experiments. We also evaluated how many candidate HLAs each method needed to account for these responses. Together, these analyses assess whether the methods can explain observed responses with a compact set of candidate HLAs, rather than directly measuring HLA restriction accuracy.

## Methods

### Model

For a fixed peptide p, let y_i_ be the number of TCRs matched to that peptide in experiment i. Let x_ik_be 1 if the donor in experiment i carries candidate HLA k and 0 otherwise. Let N_i_ be the total number of MIRA TCR matches to all other peptides in experiment i, and let NC be the mean of N_i_ across experiments. We model the count as

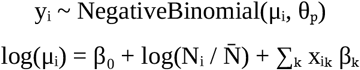

where β_k_≥ 0. We fixed the intercept β_o_ to represent low assay background rather than estimating one for each peptide. This reflects the low MIRA false-positive rate. Peptide-specific TCR counts increased with overall MIRA yield even when counts for the peptide being modeled were excluded (Supplementary Fig. 1). We therefore included log 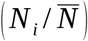, where *N*_*i*_ is the number of TCR matches to all other peptides, as an offset to account for assay yield. The coefficients β_k_are estimated jointly across candidate HLA alleles, allowing the association of each allele with peptide-specific TCR counts to be assessed while accounting for the others. The dispersion parameter θ_p_ is estimated separately for each peptide to account for variation in TCR counts.

### NetMHCpan-weighted regularization

We fit the regression with an allele-specific L1 penalty by minimizing

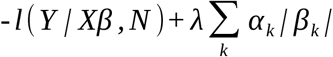

Here, α_k_is the NetMHCpan eluted-ligand percentile rank (%Rank_EL) for peptide p and HLA k. The L1 penalty encourages coefficients to become zero, leaving a small set of alleles (Tibshirani, 1996). An HLA is selected for a peptide when its fitted coefficient is positive (β_k_> 0); alleles with zero coefficients are not selected. Applying different penalty weights to different alleles follows weighted LASSO regularization (Zou, 2006).

The regularization parameter λ controls how strongly HLA coefficients are penalized, with larger values selecting fewer alleles. We tuned λ (1–20) using earlier MIRA experiments collected primarily to identify TCRs for cell therapy, separate from the held-out benchmark. Because our goal was to identify HLAs associated with peptide responses rather than predict exact TCR counts, we selected λ by minimizing the weighted misclassification rate. For each peptide, we fit the model on 80% of experiments and evaluated response predictions on the remaining 20%. An observed response required at least one peptide-specific TCR, and a predicted response required an expected TCR count greater than one. We calculated the fraction of held-out experiments misclassified for each peptide and averaged these rates across peptides, weighting each by its share of total MIRA TCR yield. The lowest weighted error occurred at λ = 4, which we fixed for the benchmark.

The penalized fit can also be interpreted as maximum-a-posteriori estimation with allele-specific Laplace priors. The important point is that NetMHCpan guides which alleles are favored, but MIRA responses determine which are retained. An allele with weaker predicted presentation can still be selected if the response data support it.

### Output scores

For each peptide, HLAResolver outputs one row for each HLA allele *k* considered by the model, including its NetMHCpan eluted-ligand rank, fitted coefficient *β*_*k*_, and a nominal coefficient *p*-value from statsmodels (Seabold and Perktold, 2010). Because this *p*-value is calculated after regularized model selection, it is descriptive rather than a formal post-selection test. We also calculate a normalized support score from the fitted coefficients:

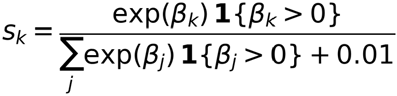

In this equation, *k* is the allele being scored, *j* indexes all candidate HLAs for that peptide, and 1 (*⋅*) is an indicator function. The constant 0.01 avoids division by zero if every coefficient is zero. The score compares candidates for the same peptide; it is not a calibrated probability.

### Held-out response prediction

To evaluate HLA assignments, we used a later MIRA dataset collected for TCR–pMHC model development, separate from the experiments used to tune HLAResolver. For each peptide, we divided HLA-typed experiments into 80% training and 20% held-out testing, keeping all TCR observations from an experiment in the same set. HLAResolver was fitted on the training data with λ = 4, fixed from the earlier tuning dataset. We included only peptides tested in panels with at least 50 valid experiments.

For all three methods, we predicted a response if a donor carried at least one HLA selected for that peptide by the respective method. We defined an observed response as at least five peptide-specific TCR matches to reduce the influence of low-count assay noise. We compared predicted and observed responses to calculate sensitivity, specificity, and positive predictive value. We also evaluated the number of candidate HLAs per peptide and per responding donor, including the fractions of responses with exactly one or no candidate HLA. Metrics were calculated separately for each peptide and averaged across peptides.

RATE was applied only to training experiments, using two-sided Fisher exact tests to assess associations between HLA carriage and peptide response. We also calculated relative frequency (RF), defined as the response frequency among allele carriers divided by the overall response frequency, requiring RF ≥ 1.3 and at least one responding carrier. We evaluated RATE and NetMHCpan at their established thresholds and at thresholds chosen to match HLAResolver’s specificity (Table 1). These comparisons assess response prediction and HLA assignment ambiguity rather than experimentally confirmed HLA restriction.

**Table 1.** Held-out response prediction and numbers of assigned HLAs, averaged across peptides. Mean candidate HLAs per peptide counts assignments before restricting them to donor genotypes; all other metrics use only alleles carried by the held-out donors.

| **Method** | **Sensitivity** | **Specificity** | **PPV** | **Mean candidate HLAs per peptide (global)** | **Mean candidate HLAs per responding donor** | **Responses with 1 candidate HLA** | **Responses with 0 candidate HLAs** |
|---|---|---|---|---|---|---|---|
| HLAResolver | 0.626 | 0.739 | 0.266 | 1.754 | 0.778 | 0.495 | 0.374 |
| NetMHCpan, specificity-matched (EL < 0.10) | 0.382 | 0.739 | 0.148 | 1.877 | 0.422 | 0.344 | 0.618 |
| NetMHCpan, strong binder (EL < 0.5) | 0.707 | 0.428 | 0.122 | 5.804 | 0.960 | 0.513 | 0.293 |
| NetMHCpan, weak binder (EL < 2.0) | 0.895 | 0.164 | 0.107 | 15.773 | 1.746 | 0.409 | 0.105 |
| RATE, specificity-matched (p < 0.215; RF ≥ 1.3) | 0.546 | 0.739 | 0.182 | 4.306 | 0.973 | 0.256 | 0.454 |
| RATE, published (p < 0.01; RF ≥ 1.3) | 0.277 | 0.932 | 0.337 | 0.636 | 0.379 | 0.186 | 0.723 |

### Linkage disequilibrium analysis

For each peptide, we calculated φ, the correlation between HLA alleles based on which training donors carried them. For each responding donor in the held-out set, we took the method’s top-ranked HLA as the lead. We counted an additional assigned HLA as a linkage disequilibrium (LD) partner if its correlation with the lead was |φ| ≥ 0.5. We excluded alleles carried by fewer than five training donors. For this analysis, RATE used p < 0.2 and RF ≥ 1.3. We calculated 95% confidence intervals by bootstrapping over peptides.

## Results

### HLA assignments and held-out response prediction

We compared HLAResolver, NetMHCpan, and RATE using held-out MIRA experiments (Fig. 2; Table 1). We evaluated how well each method predicted which donors responded to a peptide and how many candidate HLA alleles it identified. For the latter, we considered both the total number of candidates per peptide and the number carried by each responding donor. Because the true restricting HLA was unknown, these analyses measure response prediction and ambiguity in HLA assignment, not restriction accuracy.

**Figure 1.**
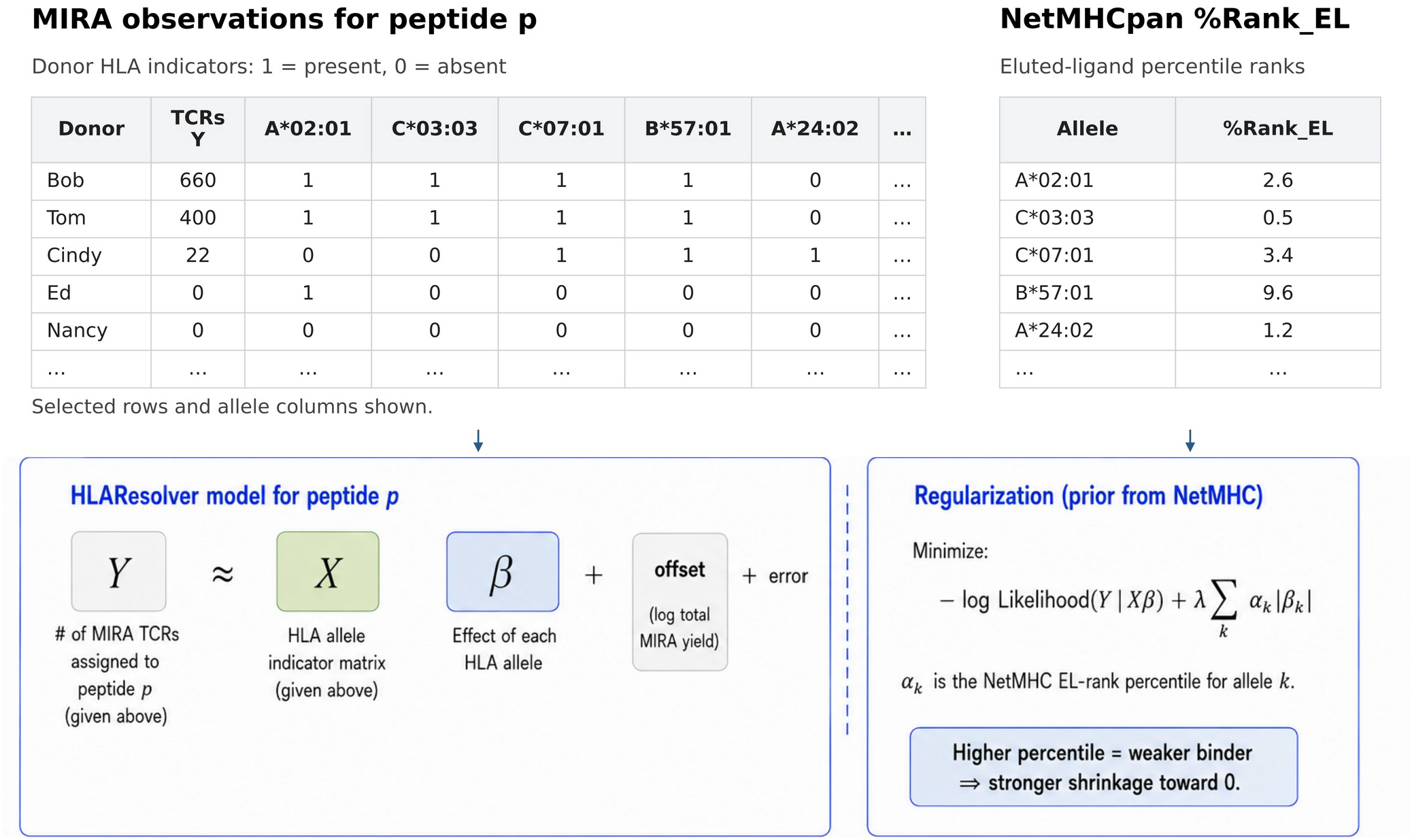
Schematic of the HLAResolver workflow. HLAResolver is run independently for each peptide. For a given peptide p, the model compares peptide-specific TCR counts across HLA-typed experiments. NetMHCpan eluted-ligand ranks guide selection of HLA alleles, and sparse regression returns a small set of candidates. The table shows only some candidate HLAs. In the model, each allele has its own presence/absence column, even if only one donor carries it. Donors who share an HLA allele have the same indicator for that allele; their other alleles can differ. The regression is illustrated schematically for intuition; the negative-binomial model and its log link are not depicted.

**Figure 2.**
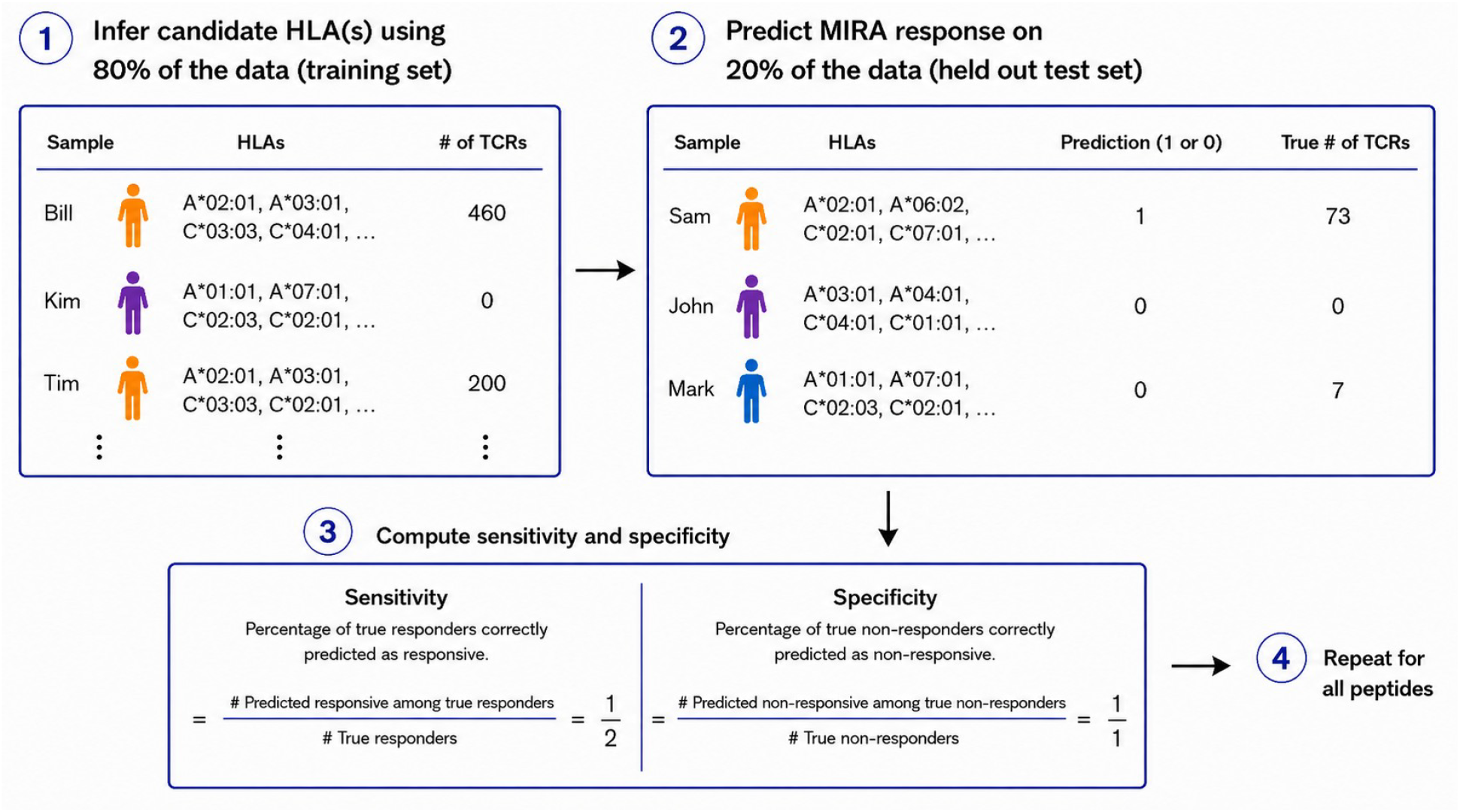
Held-out benchmark of response prediction. The 80/20 split applies to the later MIRA dataset; λ was chosen using separate, earlier experiments. For each peptide, HLAResolver was fit on 80% of HLA-typed experiments and evaluated on the remaining 20%. Each method predicted a response if a held-out donor carried at least one HLA selected for that peptide. Sensitivity and specificity describe prediction of MIRA responses, not experimentally confirmed HLA restriction.

Comparator thresholds were chosen to match HLAResolver specificity (0.739). For reference, standard NetMHCpan EL-rank cutoffs are 0.5 and 2.0 for strong and weak binders, and the published RATE cutoff is p < 0.01 with RF ≥ 1.3. Figure 3 shows the sensitivity– specificity tradeoff; Figure 4 shows how many candidates remain for each responding donor.

**Figure 3.**
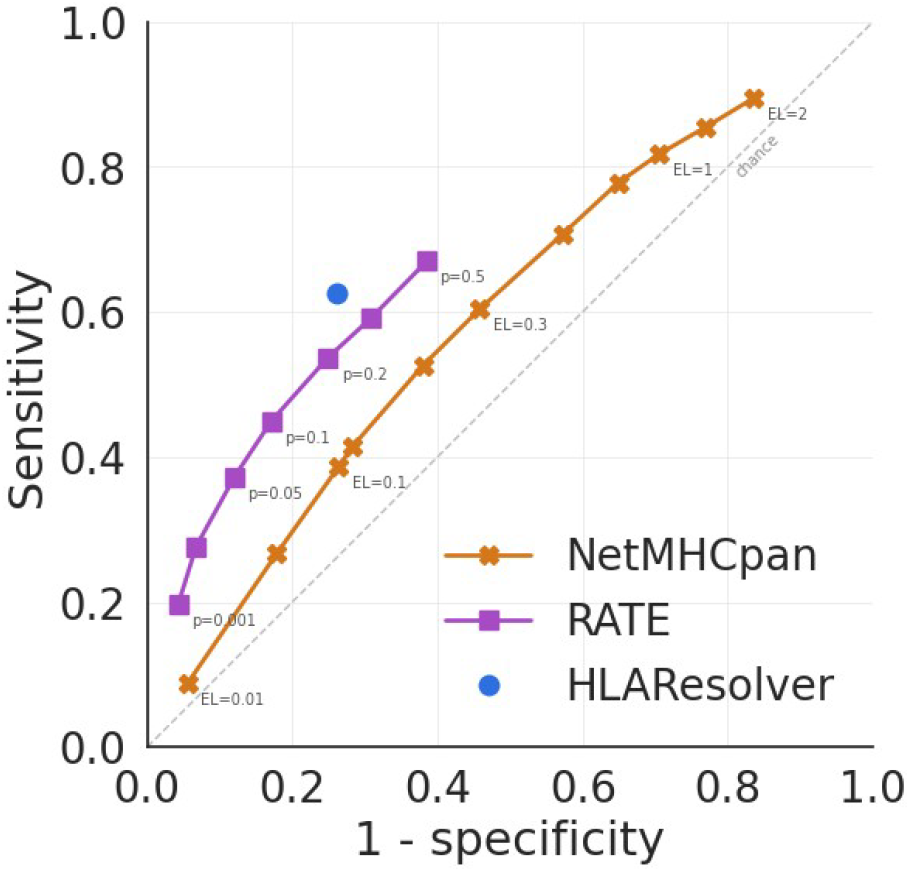
HLAResolver achieves higher sensitivity than NetMHCpan and RATE at matched specificity. Held-out MIRA response sensitivity is plotted against false-positive rate (1 − specificity). The curves show how NetMHCpan and RATE performance changes with their HLA assignment thresholds, while HLAResolver is shown as a single point. RATE used RF ≥ 1.3 throughout.

**Figure 4.**
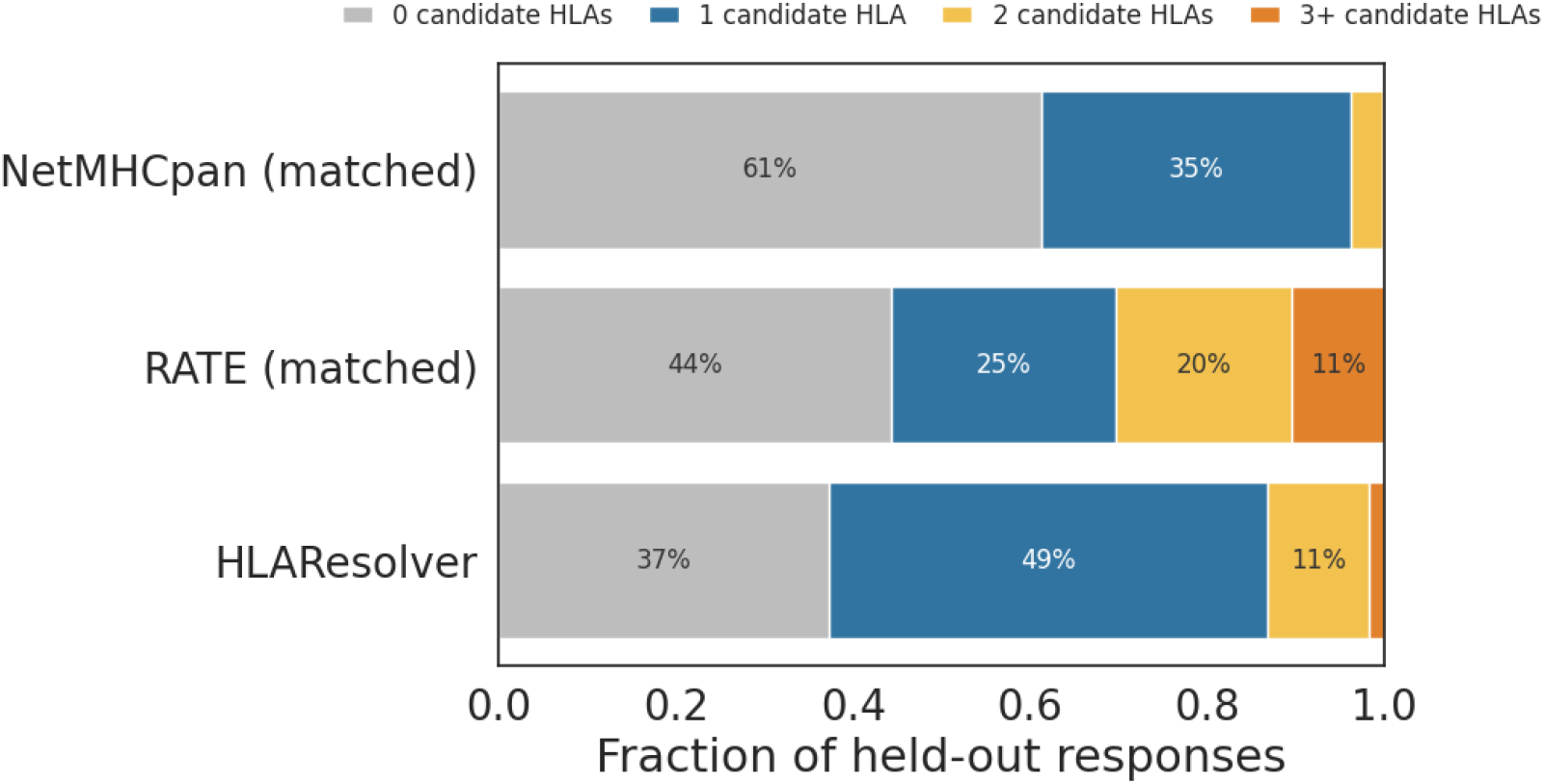
HLAResolver more often identifies a single candidate HLA for responding donors than NetMHCpan or RATE at matched specificity. For each peptide, we counted how many HLAs selected by each method were present in each responding donor’s genotype. Bars show the fractions of responding donors with zero, one, two, or three or more candidates, averaged across peptides. Table 1 reports the corresponding summary values.

At matched specificity (0.739), HLAResolver detected more observed responses than either comparator while identifying fewer candidate HLAs per peptide overall (Table 1). Its sensitivity was 0.626, compared with 0.382 for NetMHCpan and 0.546 for RATE. Thus, better response prediction did not require a larger set of candidate HLAs.

Figure 3 shows the tradeoff when the NetMHCpan and RATE thresholds are changed. At the standard NetMHCpan strong- and weak-binder thresholds, more responses were detected, but many more HLA candidates were retained and more nonresponding donors were predicted to respond. RATE at its published threshold showed the opposite pattern: higher specificity, but more missed responses. Table 1 reports the exact values for these thresholds.

### Candidate HLAs for responding donors and correlated assignments

How often can an observed MIRA response be associated with just one candidate HLA? To answer this, we considered only the candidate HLAs carried by each responding donor (Fig. 4). At matched specificity, HLAResolver identified exactly one candidate for an average of 49.5% of responding donors per peptide, more often than either comparator. NetMHCpan more often left responses without a candidate HLA, whereas RATE more often assigned multiple candidates.

NetMHCpan also assigned exactly one candidate to about half of responses at its standard strong-binder threshold. However, this came with substantially lower specificity and more than three times as many candidate HLAs per peptide overall (Table 1). The fraction of responses with one candidate should therefore be considered alongside how well the method predicts responses.

Because HLA alleles are often inherited together, a method may retain several correlated alleles even if only one presents the peptide. We therefore examined HLA calls beyond each method’s top-ranked allele. Among these additional calls, 34.2% for RATE were strongly correlated with the lead HLA (|φ| ≥ 0.5), compared with 3.8% for HLAResolver (95% CIs, 30.5–37.9% and 1.0–7.7%, respectively). This difference is consistent with HLAResolver reducing redundant assignments through joint modeling and NetMHCpan-guided selection.

### Discordance with NetMHC-designated HLA labels is concentrated in HLA-C

We next compared HLAResolver with the HLA allele originally used to select peptides for some MIRA experiments. In those experiments, peptides passed a stringent NetMHC presentation threshold for a target allele and were tested in donors carrying that allele. We call this target the NetMHC-designated HLA. Agreement with that label is not an independent validation, because both the label and HLAResolver use NetMHC predictions. Instead, the comparison shows where HLAResolver agrees with, or differs from, the intended peptide-selection target.

For most HLA-A and HLA-B alleles, HLAResolver’s top assignment matched the NetMHC-designated HLA (Fig. 5). Agreement was nearly complete for A*11:01, A*68:01, B*57:01, B*35:01, and B*15:01. The largest disagreements were for HLA-C, especially C*16:01, where the model often favored another plausible HLA-C allele. HLA-C alleles have more similar and overlapping peptide-binding motifs than many HLA-A and HLA-B alleles, which could contribute to these disagreements (Sarkizova et al., 2020).

**Figure 5.**
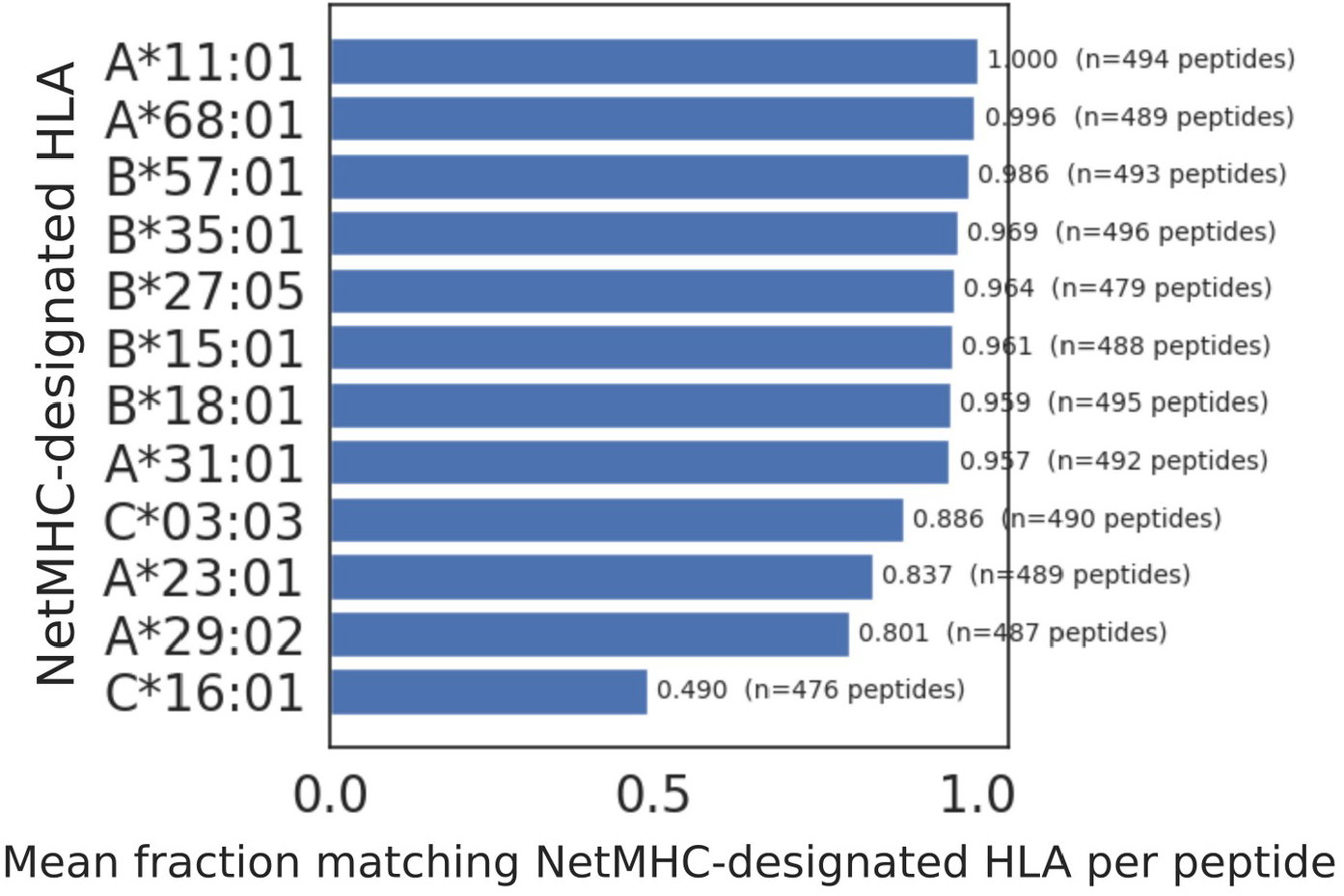
HLAResolver generally agrees with NetMHC-designated HLAs for HLA-A and HLA-B, but less often for HLA-C. Bars show the fraction of peptides for which HLAResolver’s top assignment matched the HLA originally used for peptide selection. These designated HLAs are based on NetMHC predictions rather than experimentally confirmed restrictions.

The concentration of disagreements in HLA-C suggests that assigning a single presenting allele may be more challenging for these peptides, potentially because of overlapping binding preferences among HLA-C alleles. These cases warrant greater caution when assigning HLA restrictions for TCR–pMHC model training, particularly when the available evidence supports multiple plausible alleles.

## Discussion

MIRA links TCRs to peptides but does not reveal which HLA presents each peptide. HLAResolver addresses this gap by combining NetMHCpan predictions of peptide presentation with MIRA response patterns across HLA-typed donors. By evaluating candidate alleles jointly, the model uses both sources of evidence to select a compact set of HLAs while reducing redundant assignments among alleles correlated through linkage disequilibrium. The resulting annotations narrow the possible presenting HLAs for TCR–pMHC model training.

The held-out benchmark shows that HLAResolver’s assignments predict response patterns in new experiments. At matched specificity, HLAResolver accounted for more observed responses while retaining a compact set of candidate HLAs per peptide. This advantage may reflect the complementary strengths of the two sources of evidence: NetMHCpan prioritizes alleles likely to present a peptide, while MIRA response patterns provide statistical support for HLA associations.

Additional assignments were less often strongly correlated with the lead HLA for HLAResolver than for RATE (3.8% versus 34.2%), consistent with reduced redundancy. More permissive NetMHCpan thresholds detected additional responses but assigned many more HLAs, whereas stricter RATE thresholds missed more responses. RATE was evaluated only in panels with at least 50 valid experiments, so its performance in smaller panels remains unknown.

The method has several limitations. First, MIRA does not establish the true restricting HLA, and experimental validation is not available at this scale. Our benchmark therefore measures how well the inferred HLAs explain observed responses, not whether they correctly identify the restricting allele. Second, sparse regression may miss additional valid presenting HLAs when a peptide is presented by more than one allele. Third, when multiple candidate HLAs remain within a donor, HLAResolver cannot determine which allele restricts each individual peptide-responsive TCR. Relative support scores should not be interpreted as the proportions of TCRs restricted by each allele. Fourth, highly correlated alleles may remain difficult to distinguish when response patterns and NetMHCpan predictions are similar. Finally, MIRA TCR counts reflect repertoire sampling, assay yield, immunodominance, and other factors besides peptide presentation.

HLAResolver combines predicted peptide presentation with MIRA response patterns to identify compact sets of candidate HLAs for TCR–peptide associations. By narrowing the possible presenting HLAs, it makes large-scale MIRA data more useful for training TCR–pMHC models.

