## Supplementary Figure 1 for "HLAResolver: A statistical framework for resolving HLA ambiguity in T-cell response data"

### HLAResolver — Supplementary information

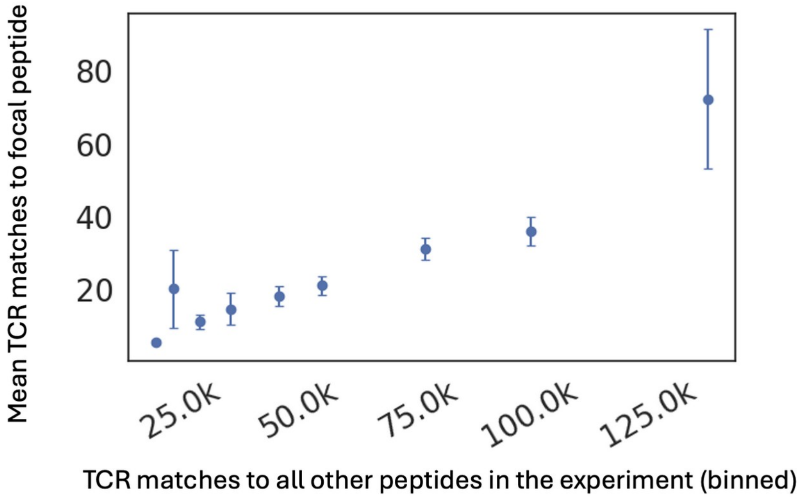


**Supplementary Figure 1. Relationship between focal-peptide and experiment-level MIRA yield.** Focal yield counts TCRs matched to the focal peptide; off-target yield counts matches to other peptides in the same experiment. Points show off-target-yield bin means ± SE. Their positive relationship motivated the offset $\log\left(N_i/\overline{N}\right)$ for experiment-level MIRA yield.
